# Hepatic Lipid Droplet-Associated Proteome Changes Distinguish Dietary-Induced Fatty Liver from Insulin Resistance in Male Mice

**DOI:** 10.1101/2023.03.09.531813

**Authors:** Andries Van Woerkom, Dylan J Harney, Shilpa R. Nagarajan, Mariam F. Hakeem-Sanni, Jinfeng Lin, Matthew Hooke, Tamara Pulpitel, Gregory J Cooney, Mark Larance, Darren N. Saunders, Amanda E Brandon, Andrew J. Hoy

**Affiliations:** School of Medical Sciences, Charles Perkins Centre, Faculty of Medicine and Health, University of Sydney, NSW 2006, Australia.; School of Life and Environmental Sciences, Charles Perkins Centre, Faculty of Science, The University of Sydney, NSW 2006, Australia

## Abstract

Fatty liver is characterised by the expansion of lipid droplets and is associated with the development of many metabolic diseases, including insulin resistance, dyslipidaemia and cardiovascular disease. We assessed the morphology of hepatic lipid droplets and performed quantitative proteomics in lean, glucose-tolerant mice compared to high-fat diet (HFD) fed mice that displayed hepatic steatosis and glucose intolerance as well as high-starch diet (HStD) fed mice who exhibited similar levels of hepatic steatosis but remained glucose tolerant. Both HFD and HStD-fed mice had more and larger lipid droplets than Chow-fed animals. We observed striking differences in liver lipid droplet proteomes of HFD and HStD-fed mice compared to Chow-fed mice, with fewer differences between HFD and HStD. Taking advantage of our diet strategy, we identified a fatty liver lipid droplet proteome consisting of proteins common in HFD- and HStD-fed mice. Likewise, a proteome associated with glucose tolerance that included proteins common in Chow and HStD but not HFD-fed mice was identified. Notably, glucose intolerance was associated with changes in the ratio of adipose triglyceride lipase (ATGL) to perilipin 5 (PLIN5) in the lipid droplet proteome, suggesting dysregulation of neutral lipid homeostasis in glucose-intolerant fatty liver, which supports bioactive lipid synthesis and impairs hepatic insulin action. We conclude that our novel dietary approach uncouples ectopic lipid burden from insulin resistance-associated changes in the hepatic lipid droplet proteome.

## INTRODUCTION

Fatty liver is an early and defining feature of a range of liver diseases, including non alcoholic fatty liver disease (NAFLD [1]), alcoholic liver disease [2], hepatitis C [3] and HIV [4]. If not addressed, fatty liver can progress to steatohepatitis, cirrhosis and carcinoma; in fact, enhanced synthesis of lipids, including fatty acids, glycerolipids and sphingolipids, is essential for mTORC2- mediated hepatocellular carcinoma [5]. It has been estimated that ∼25% of the world’s population is currently thought to have NAFLD, and this is predicted to increase significantly, leading to increased death due to liver-related pathologies [6]. The increasing prevalence of fatty liver reflects the increasing prevalence of other non-communicable cardiometabolic diseases, such as type 2 diabetes, obesity, and cardiovascular disease [7].

Fatty liver is highly associated with obesity [8]; however, the clinical manifestations of obesity are heterogeneous and complex, as is the prevalence of associated disease [9–12]. For example, it has been estimated that one-third of patients with obesity are metabolically healthy; the remaining being ‘obese-metabolically unhealthy’ [13], highlighting metabolic diversity within a population defined as obese by BMI. Similarly, it has been estimated that ∼5-8% of patients with NAFLD in Western countries are considered lean, whereas ∼20% of the Asian population has lean NAFLD [6]. Whilst currently there are no universally accepted criteria for identifying metabolically-(un)healthy individuals, generally, it includes a combination of adiposity, insulin sensitivity, inflammation and circulating glucose and lipids [14, 15]. Importantly, the incidence of NAFLD is strongly associated with being overweight and obesity, even in well-defined metabolically healthy men and women [16].

NAFLD is defined as intrahepatic triacylglycerol content greater than 5%, which histologically equates to an increase in the number and size of intracellular lipid droplets [1]. Lipid droplets are dynamic organelles that store neutral lipids like triacylglycerol (TG) and cholesteryl esters and are coated by a large number of proteins, some of which are known to be involved in the incorporation or release of lipids from the droplets [17]. The accumulation of triacylglycerols results from an imbalance between the uptake of extracellular lipids, *de novo* synthesis of fatty acids, oxidation and release of TG-VLDL [18]. In rodent models, lipid accumulation in the liver is an early event in the development of high-fat diet-induced insulin resistance and obesity [19, 20]. However, even in animal models, little knowledge exists regarding how lipid storage in liver cells is dysregulated in insulin resistance. Moreover, there is little understanding of the relationship between liver lipid storage and insulin action in a setting of metabolically healthy obesity.

To date, the lipid droplet proteome has been defined for a diverse array of organisms and cell and tissue types (see review [17]), including in rodent liver and in models of obesity and NAFLD [21–26]. It is important to highlight that these studies predominantly compare two groups, for example, between Chow (control, low-fat diet) and high-fat diet-fed animals or fasting and fasting-re-fed states, which result in multiple differences between two groups, such as adiposity, tissue and circulating lipid levels, circulating hormone levels, and immune status. This binary normal vs obese framework fails to capture the metabolic diversity of the obese population (i.e. ‘obese-metabolically unhealthy’ vs ‘obese-metabolically healthy’ etc. [13, 27]) as well as being unable to separate the role of lipid accumulation from hyperinsulinaemia and altered immune status. Recently, we showed that feeding mice a diet high in starch induced obesity and lipid accumulation in liver and skeletal muscle to levels similar to mice fed a high-fat diet, but that the high-starch diet group retained glucose tolerance and insulin sensitivity when compared to mice fed a high-fat diet [28]. This approach provides a powerful platform to identify the molecular mechanisms that result in the “safe” and “unsafe” storage of excess lipids and its links to insulin sensitivity, analogous to the ‘Athlete’s Paradox’ where highly insulin-sensitive, endurance-trained athletes have skeletal muscle lipid levels similar to that observed in insulin-resistant obese and type 2 diabetes subjects [29]. In this study, we aimed to deploy this model – using mice fed either a high-fat diet (HFD), a high-starch diet (HStD), or Chow control - to determine changes in liver lipid droplet morphology and proteome associated with glucose tolerance from changes linked to liver lipid content.

## RESULTS

### High-fat and high-starch diet feeding leads to increased adiposity and liver triacylglycerol content but differs in glucose tolerance

In line with our previous observations [28], mice fed either an HFD or HStD had increased energy intake (Figure 1B) and increased body weight (Figure 1C) compared to mice fed Chow. Further, mice fed either HFD or HStD had greater total fat mass (Figure 1D) and epidydimal and subcutaneous fat pad mass (Figure 1E), as well as liver TG content (Figure 1F) compared to Chow-fed controls.

**Figure 1:**
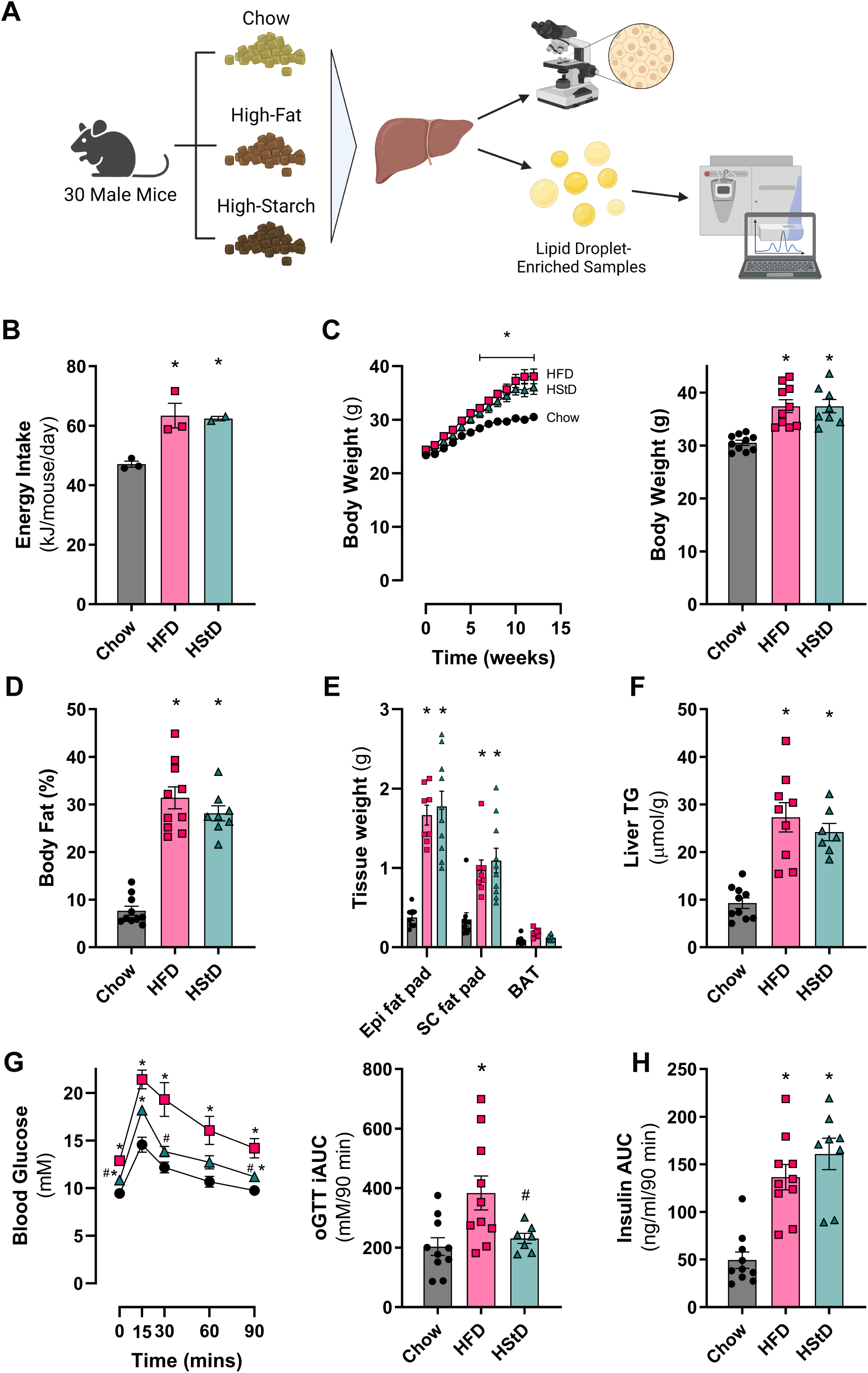
HStD and HFD similarly increases food intake, body weight, adiposity and liver triacylglycerol levels but HStD retain glucose tolerance compared to HFD. (A) Experimental design of the morphometric and proteomic analyses of liver samples enriched for lipid droplets from mice fed Chow diet, high-fat diet (HFD) or high-starch diet (HStD). Created with BioRender.com (B) Average daily energy intake, (C) body weight and end point body weight, (D) body fat mass, (E) tissue weights, (F) liver triacylglycerols (TG), (G) blood glucose levels and incremental area under the curve (iAUC) for the oral glucose tolerance test after 12 weeks of feeding, (H) incremental area under the curve of insulin levels for the oral glucose tolerance test after 12 weeks of feeding. Data are presented as mean ± SEM; (B) n=3 for Chow and HFD, n=2 for HStD; (C-H) n=10 for Chow, n=9 for HFD, n=8 for HStD. * *P*≤ 0.05 vs. Chow; # *P*≤ 0.05 vs. HFD by One-way ANOVA (B, C right panel, D-F, G right panel, H) or Two-way ANOVA (C left panel & G left panel) followed by Tukey’s Multiple Comparisons test.

Despite the similarity in body weight, adiposity and liver lipid content, mice fed HStD remained glucose tolerant, whereas mice fed HFD were glucose intolerant (Figure 1G). The difference in glucose tolerance was not driven by differences in insulin release (Figure 1H). Collectively, these data demonstrate that HFD feeding led to increased adiposity, hepatic steatosis and glucose intolerance, whereas HStD feeding resulted in similar levels of adiposity and hepatic steatosis but did not induce glucose intolerance. These observations are consistent with our previous report, where HStD feeding resulted in no evidence of hepatic insulin resistance, as determined using the gold-standard hyperinsulinaemic-euglycaemic clamp technique [28].

### Hepatic lipid droplet morphology is similar between mice fed HFD and HStD

The differences in whole-body glucose metabolism between mice fed HFD and HStD, despite matched liver lipid levels, suggest that mice fed HStD can safely store excess lipids in cytosolic lipid droplets, whereas mice fed HFD do not. As such, we determined whether differences in glucose tolerance were associated with liver lipid droplet morphology. Consistent with the biochemical measure of TG (Figure 1F), an increased number and size of hepatic lipid droplets were observed in mice fed an HFD or HStD compared to mice fed a Chow diet (Figure 2). However, there was no difference in the number and size of liver lipid droplets between HFD-fed mice and those provided with an HStD (Figure 2). This suggests that the differences in whole-body glucose metabolism between mice fed an HFD and HStD were not associated with changes in the morphology of liver lipid droplets, despite both groups exhibiting liver steatosis.

**Figure 2:**
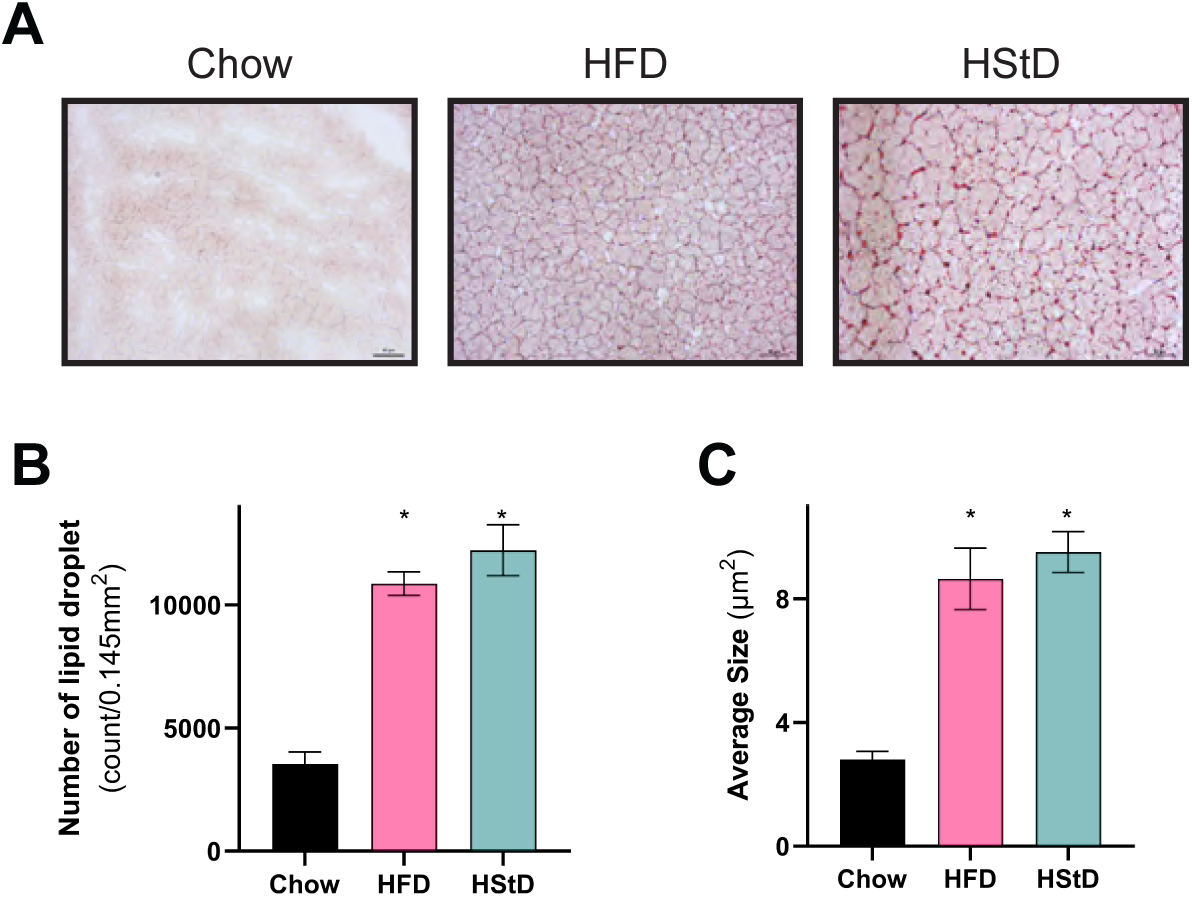
HStD and HFD similarly increases liver lipid droplet number and size compared with Chow. (A) Representative Oil Red-O stained mouse livers from mice fed Chow diet, high-fat diet (HFD) or high-starch diet (HStD) for 12 weeks. (B) Lipid droplet number per area and (C) size. Data are presented as mean ± SEM; Up to 4 regions of interest quantified per mouse. n=10 for Chow, n=9 for HFD, n=8 for HStD. * *P*≤ 0.05 vs. Chow by One-way ANOVA followed by Tukey’s Multiple Comparisons test.

### High-fat diet induces a distinct liver lipid droplet proteome compared with a high-starch diet

We propose that the development of glucose intolerance is associated with molecular events that regulate the storage of excess lipids. Hence, we next quantified the proteomes of liver lipid droplets in mice fed our three diets. Lipid droplets were enriched by gently homogenising fresh liver and sucrose gradient centrifugation [30]. Enrichment was confirmed by immunoblot detection of established markers (Figure 3A), and samples were delipidated prior to mass spectrometry analyses. We identified 1968 proteins in LD-enriched samples from Chow-fed mice that met our inclusion criteria of a minimum of two peptides for each protein in at least 7 of 10 mice for Chow (Table S2). Similarly, we identified 2108 proteins in LD-enriched samples in 7 of 9 mice for HFD from HFD-fed mice and 2030 proteins in 7 of 8 mice for HStD-fed mice (Table S3). Of these identified proteins, 1823 were common to all three groups (Table S4; Figure 3 - supplement S3A). The median Pearson correlation coefficient for the LD-associated proteome for each diet was 0.928 for Chow, 0.965 for HFD, and 0.927 for HStD (Figure 3 - supplement S3B).

**Figure 3:**
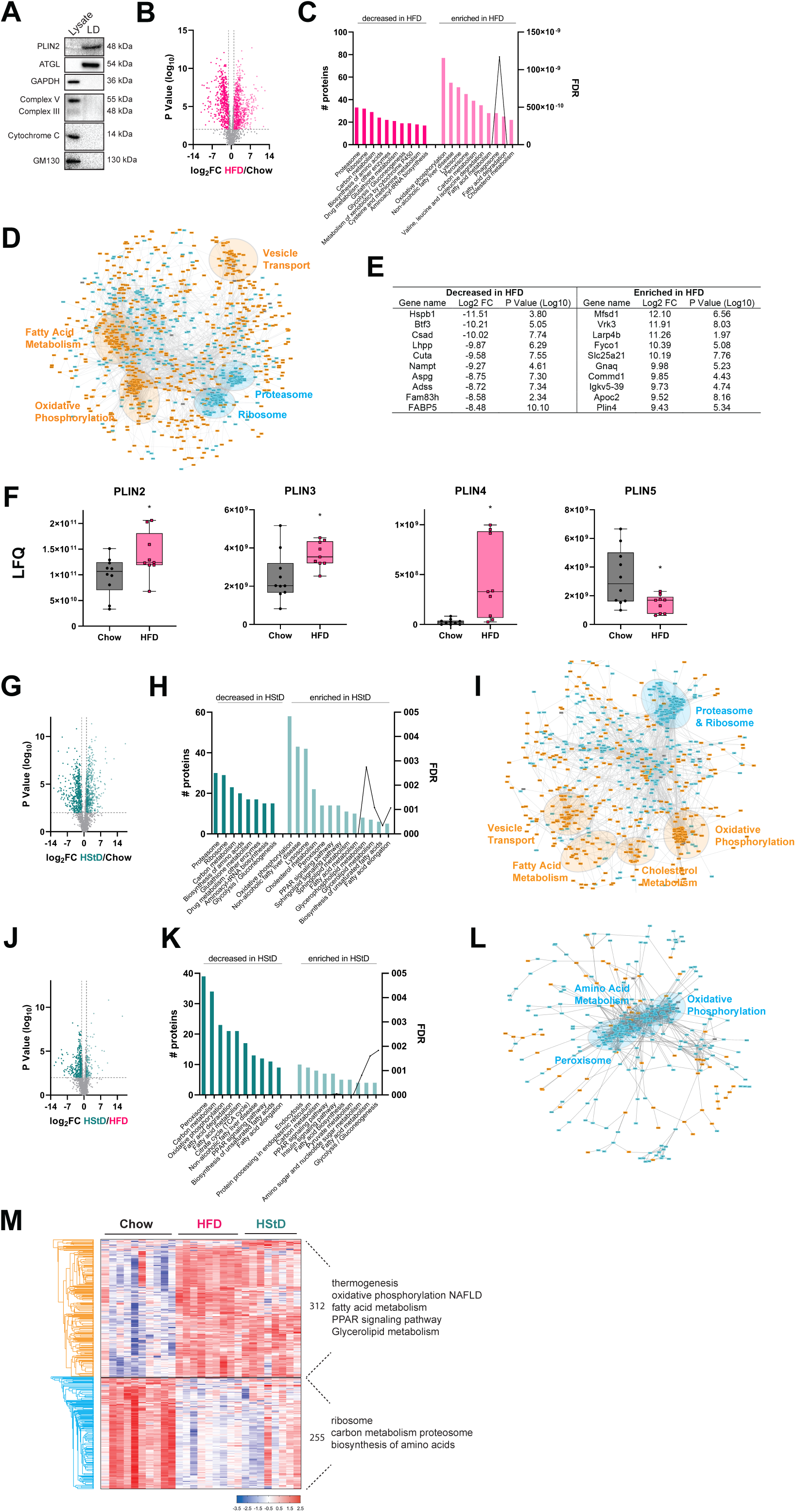
The proteome of liver lipid droplets is modified by HFD and HStD feeding. (A) Representative immunoblots of protein markers of organelles including lipid droplets (Plin2, ATGL), cytosol (GAPDH), mitochondria (protein subunits in the mitochondrial complexes (complex III-Core protein 2 and complex V-alpha subunit, cytochrome C), and golgi (GM130). (B) Volcano plot of lipid droplet-associated proteins in response to high-fat diet (HFD) feeding compared to Chow diet feeding. (C) Ontology analysis and (D) STRING analysis of significantly changed lipid droplet associated proteins in response to HFD feeding compared to Chow diet feeding. (E) List of proteins most significantly altered in abundance in the lipid droplet proteome following 12 weeks of HFD feeding. (F) Box and whisker plots for specific proteins of interest. Each point represents protein abundance from an individual mouse. (G) Volcano plot of lipid droplet-associated proteins in response to high-starch diet (HStD) feeding compared to Chow diet feeding. (H) Ontology analysis and (I) STRING analysis of significantly changed lipid droplet associated proteins in response to HStD feeding compared to Chow diet feeding. (J) Volcano plot of lipid droplet-associated proteins in response to HStD feeding compared to HFD diet feeding. (K) Ontology analysis and (L) STRING analysis of significantly changed lipid droplet associated proteins in response to HStD feeding compared to HFD diet feeding. (M) Hierarchical clustering of label-free quantitation (LFQ) intensities of 567 significantly changed proteins (ANOVA, FDR 0.05) in the LD-enriched proteome revealed two clusters related to changes in response to HFD and HStD feeding. Numbers of proteins and selected enriched KEGG pathways (Fisher’s exact test, FDR 0.1) are indicated for marked clusters. n=10 for Chow, n=9 for HFD, n=8 for HStD. Data in (F) are presented as box and whisker plots: median, interquartile range and error bars representing Min to Max. * P≤ 0.05 vs. Chow by t-test. LFQ, label-free quantification

Quantitative analysis of changes in LD protein abundance was next performed using LFQ values. From this analysis, we identified 1349 proteins with altered abundance in the LD-enriched samples from livers of HFD-fed animals compared to Chow, indicating that a significant proportion (74%) of the LD proteome is responsive to HFD feeding (Figure 3B, Table S5). Of these, 806 proteins in the LD-enriched fraction had increased abundance with HFD, whereas 543 had relatively lower abundance (Figure 3B). KEGG pathway analysis of enriched proteins identified enrichment of components of NAFLD, fatty acid metabolism and glycerolipid metabolism, whilst those proteins decreased in abundance were involved in carbon metabolism among other pathways (Figure 3C). STRING analysis of protein-protein interaction (PPI) networks within the set of significantly changed LD-associated proteins (n = 817) in animals fed an HFD identified enrichment for components of fatty acid metabolism, vehicle transport, and oxidative phosphorylation and decreased proteasome and ribosome proteins (Figure 3D).

Notably, several of the top 10 proteins that were most increased or decreased in liver LD-enriched fractions of HFD-fed mice included proteins that have been reported to influence liver lipid metabolism, including COMM domain-containing protein 1 (Commd1) [31], nicotinamide phosphoribosyltransferase (Nampt) [32], fatty acid binding protein 5 (FABP5) [32], and PLIN4 [33] (Figure 3E). Furthermore, the protein levels of many known LD-associated regulators of TG lipolysis were, as expected, significantly altered in response to HFD feeding compared to Chow-fed mice, including PLIN2, PLIN3, PLIN4, PLIN5 (Figure 3F), hormone sensitive lipase (HSL; FC = 1.73, p=0.016), G0S2 (FC = 1.58, p=0.019), and ABHD5 (also known as CGI58; FC = -4.4, p<0.0001), with the notable exception being ATGL (FC = 1.22, p=0.19).

Our novel HStD induces obesity and fatty liver in C57BL/6Arc mice yet does not lead to the development of glucose intolerance or insulin resistance, compared to HFD-fed mice (Figure 1) [28]. We next analysed LD-enriched samples from mice fed the HStD to those from Chow-fed mice to identify changes in protein levels in another fatty liver model but one that retains insulin sensitivity. From this analysis, we identified 1021 (56% proteins) of the proteome was differentially regulated in the LD-enriched fractions of the livers of mice an HStD compared to mice (Table S6). Specifically, there were 531 proteins in the LD-enriched fraction that were increased, and 490 decreased in abundance in response to HStD feeding (Figure 3G). Unsurprisingly for mice with fatty liver, the enriched LD-associated proteins were primarily involved in cholesterol, and fatty acid metabolism, glycerolipid metabolism and fatty acid elongation and desaturation (Figure 3H), which is distinct from HFD-fed mice (Figure 3C). The increased abundance of proteins involved in fatty acid synthesis and modification is consistent with the notion that HStD-fed mice have to synthesise fatty acids whereas HFD-fed mice have abundant access to fat from the diet [28]. Those proteins decreased in abundance in HStD-fed mice were involved in carbon, glucose and amino acid metabolism (Figure 3H). PPI analysis of the significantly altered proteins identified clusters of proteins involved in fatty acid and cholesterol metabolism, oxidative phosphorylation, and decreased proteasome and ribosome proteins (Figure 3I). Finally, we also identified 467 (26% proteins) significantly altered proteins in LD-enriched fractions of livers from HStD-fed mice compared to HFD mice (Table S7), two groups with equal levels of liver TGs (Figure 1F). Specifically, 116 proteins in the LD-enriched fraction were increased, and 351 decreased in abundance in response to HStD (Figure 3J). The enriched LD-associated proteins in HStD-fed mice were primarily involved in fatty acid biosynthesis and metabolism, whereas those proteins that decreased in abundance were listed in the NAFLD and fatty acid metabolism KEGG pathways (Figure 3K). We also observed pathways including carbon and fatty acid metabolism and PPAR signalling were increased and decreased, which can be explained by members of these pathways be differentially affected by HFD and HStD feeding. Significantly altered proteins underwent PPI analysis that identified clusters of proteins involved in amino acid metabolism, oxidative phosphorylation and peroxisome that were downregulated in the HStD compared to HFD (Figure 3L). Most interesting is the enrichment of proteins involved in oxidative phosphorylation in LD-enriched sampled from HFD-fed mice compared to HStD-fed mice, which suggests that there are dietary specific effects on LD-mitochondrial interactions and thereby mitochondrial function [34, 35]. Of note, there were striking differences in the patterns observed when comparing the STRING PPI analysis reported in Figures 3D, I & L. These differences highlight a greater influence of the lipid levels in mouse liver (i.e. Figures 3D & I) compared to glucose tolerance (Figure 3L) on LD-associated proteome changes.

We next assessed the proteome data for all samples by ANOVA and identified 567 significantly changed proteins (Table S8). Using hierarchical cluster analysis, we identified two clear clusters, one containing 312 proteins that are primarily involved in thermogenesis, oxidative phosphorylation, NAFLD, and fatty acid metabolism. In contrast, the second cluster had 255 proteins involved in the ribosome, carbon metabolism, proteasome, and biosynthesis of amino acids (Figure 3M). It was not overly surprising that we only identified two clusters in our ANOVA analysis since the HStD induced a phenotype that exhibits similar traits to both Chow-fed animals and HFD-fed animals (Figure 1). Collectively, we show that the proteome of liver lipid droplets of C57BL/6Arc mice was altered in response to HFD and HStD feeding and that there were clear differences between these groups despite being equally obese and exhibiting similar levels of liver steatosis.

### The fatty liver lipid droplet proteome

Our dietary approach provides a unique opportunity to uncouple the changes in the LD proteome of fatty liver from those changes associated with glucose tolerance and insulin sensitivity. Firstly we filtered our data through a biologically relevant scenario to identify proteins whose abundance correlated with liver TG levels (Figure 4A); specifically, proteins that were increased/decreased in HFD compared to Chow (p ≤ 0.05) and increased/decreased in HStD compared to Chow (p ≤ 0.05) but not different between HStD and HFD (p ≥ 0.05). Using this approach, we identified 283 proteins that were increased and 285 proteins that were decreased in this scenario (Figure 4B, Table S9). KEGG pathway analysis of these proteins identified the enrichment of components of NAFLD and carbon, amino acid and glycerolipid metabolism (Figure 4C). Strikingly, many proteins known to be involved in liver lipid metabolism were also identified as being increased in the LD-enriched fractions of fatty liver, including PLIN4, monoacylglycerol lipase (MGLL) and mitoguardin 2 (MIGA2) (Figure 4D), whilst fatty acid binding protein 1 (FABP1) and acyl-CoA-binding protein (ACBP_Dbi) were decreased (Figure 4E). Other members of the PLIN family, PLIN2 and PLIN3, did not meet our criteria, but it is important to note that PLIN2 was increased in HStD samples compared to Chow but not significantly different between Chow and HFD (p=0.15 by One-Way ANOVA; Figure 4F). PLIN3 was not significantly increased in HFD compared to Chow (p=0.12 by One-Way ANOVA) but was increased in HStD compared to Chow and further enriched in LD-associated samples from HStD samples compared to HFD (by One-Way ANOVA; Figure 4F).

**Figure 4:**
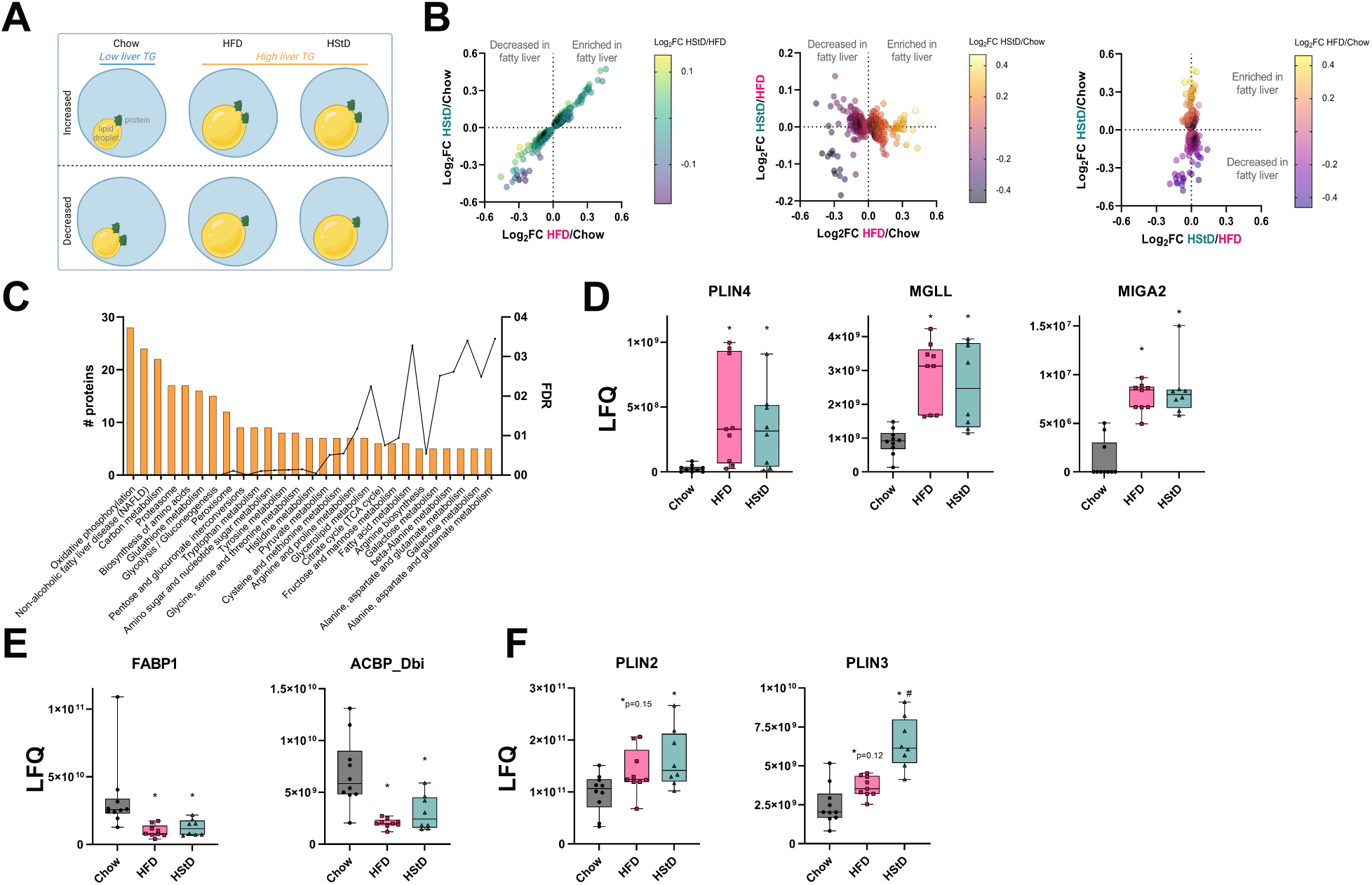
The fatty liver lipid droplet-associated proteome. (A) Biological scenario used to identify the fatty liver lipid droplet-associated proteome; increased/decreased in HFD compared to Chow (p ≤0.05) and increased/decreased in HStD compared to Chow (p ≤0.05) but not different between HStD and HFD (p≤ 0.05). Created with BioRender.com (B) The relationship of the Log2FC of proteins that were identified from the biological scenario data curation to identify fatty liver-associated changes in the lipid droplet proteome. (C) Ontology analysis of significantly changed lipid droplet-associated proteins that associate with fatty liver. Box-and-whisker plots for known lipid metabolism proteins that were (D) enriched and (E) reduced in the lipid droplet- associated proteins that associate with fatty liver. (F) Box-and-whisker plots for PLIN2 and PLIN3 abundance in the LD-enriched sampled from mice fed Chow, HFD or HStD for 12 weeks. Each point represents the protein abundance in lipid droplet-enriched fractions for an individual mouse. n=10 for Chow, n=9 for HFD, n=8 for HStD. Data in (D)-(F) are presented as box and whisker plots: median, interquartile range and error bars representing Min to Max. * P≤ 0.05 vs. Chow; # *P* ≤ 0.05 vs. HFD by One-way ANOVA followed by Tukey’s Multiple Comparisons test. LFQ, label-free quantification.

Overall, we have defined the liver LD-associated proteome common to obese mice fed HFD and HStD, independent of the differences in glucose tolerance and insulin sensitivity between these groups.

### The glucose-intolerant liver lipid droplet proteome

Similar to our approach to identifying changes in protein levels of the LD-enriched fraction of liver from mice reported above, we filtered our data through another biologically relevant scenario to identify LD-associated proteins whose abundance correlated with glucose tolerance and insulin sensitivity (Figure 5A); specifically, proteins that were increased/decreased in HFD compared to Chow (p ≤ 0.05) and increased/decreased in HFD compared to HStD (p ≤ 0.05) but not different between HStD and Chow (p ≥ 0.05). Using this approach, we identified 61 proteins that were increased and only 19 proteins that were decreased in this scenario (Figure 5B; Table S10). KEGG pathway analysis of these proteins identified the enrichment of components of fatty acid metabolism, degradation, elongation and desaturation, as well as PPAR signalling (Figure 5C). Many key proteins involved in liver lipid droplet homeostasis were identified as being decreased in the LD-enriched fractions of the liver of insulin-resistant mice, including PLIN5, ABHD5, fatty acid binding protein 4 (FABP4) and acyl-CoA synthetase long-chain family member 4 (ACSL4) (Figure 5D) whilst acetyl-CoA acetyltransferase 1 (ACAT1), which catalyses the esterification of cholesterol, was increased (Figure 5E).

**Figure 5:**
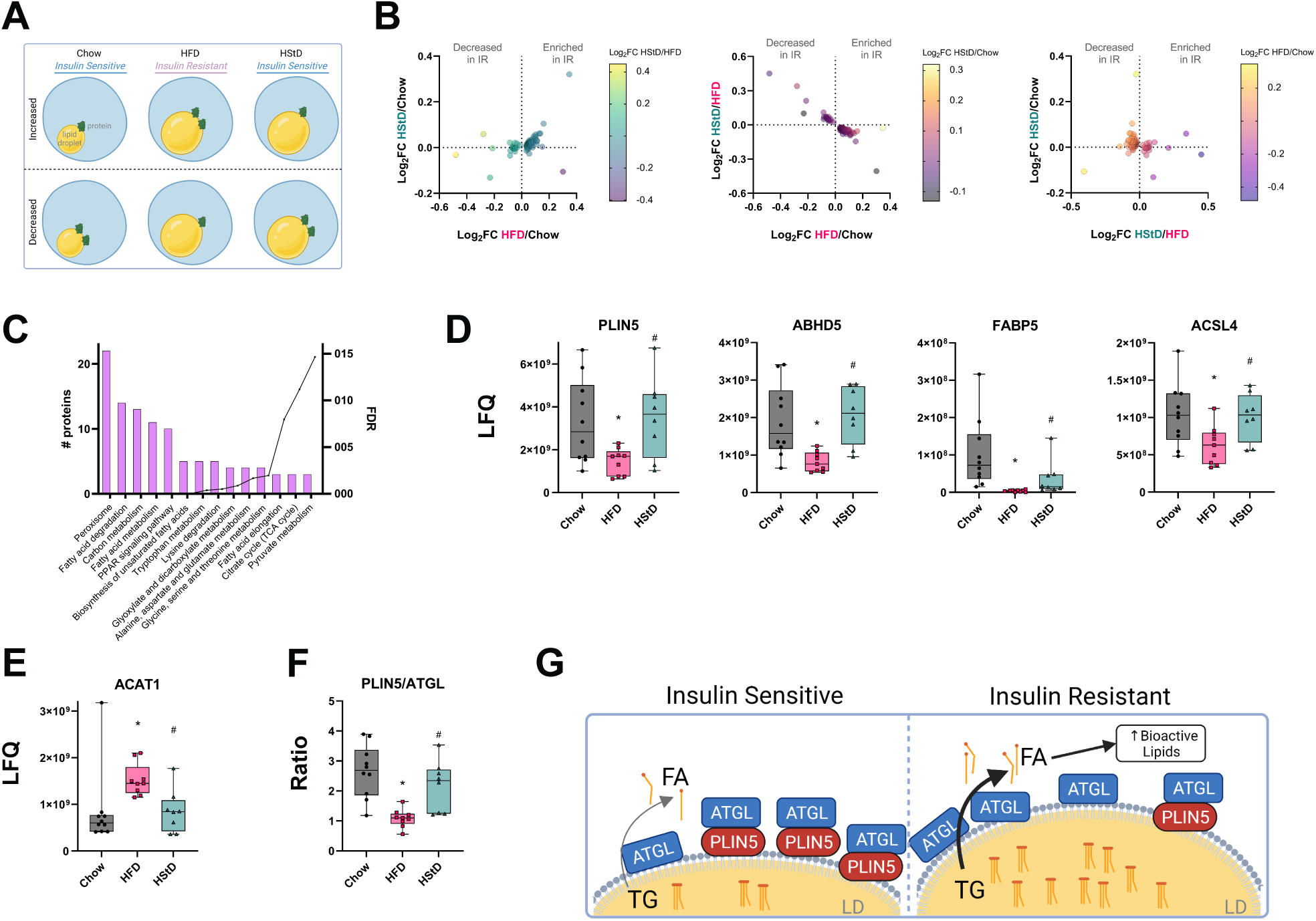
The liver lipid droplet-associated proteome that associates with glucose tolerance and insulin sensitivity. (A) Biological scenario used to identify the liver lipid droplet-associated proteome that associates with insulin resistance; increased/decreased in HFD compared to Chow (p≤0.05) and increased/decreased in HFD compared to HStD (p≤0.05) but not different between HStD and Chow (p ≥ 0.05). Created with BioRender.com (B) The relationship of the Log2FC of proteins that were identified from the biological scenario data curation to identify insulin resistance- associated changes in the lipid droplet proteome. (C) Ontology analysis of significantly changed liver lipid droplet-associated proteins that associate with glucose tolerance and insulin sensitivity. Box-and-whisker plots for known lipid metabolism proteins that were (D) enriched or (E) decreased in the lipid droplet-associated proteins that relate with glucose tolerance and insulin sensitivity. (F) The PLIN5/ATGL ratio of LFQ data. Each point represents the protein abundance in lipid droplet- enriched fractions for an individual mouse. (I) Proposed model of changes in the liver lipid droplet-associated protein that aligns with glucose tolerance and insulin sensitivity. n=10 for Chow, n=9 for HFD, n=8 for HStD. Data in (D)-(F) are presented as box and whisker plots: median, interquartile range and error bars representing Min to Max. * P≤0.05 vs. Chow; # *P*≤0.05 vs. HFD by One-way ANOVA followed by Tukey’s Multiple Comparisons test. LFQ, label-free quantification.

Of most interest were the changes in PLIN5, which acts as a gatekeeper for access to lipid droplet- contained substrates [36]. So we examined other known lipid metabolism proteins to put into context the changes in PLIN5. From this, we observed no changes in the protein levels of DGAT2, ATGL and G0S2 between all three groups and a broad range of differences between groups for other proteins (Figure 5 - supplement S5). ATGL activity is influenced by protein-protein interactions with its coactivators and co-suppressor [37–39], phosphorylation [40, 41] and translocation [42, 43], and access to its LD-contained substrates [36]. Since we observed no difference in ATGL protein levels between groups (Figure 5 - supplement S5), we proposed that in the insulin-resistant setting, there may be dysregulated ATGL activity at the lipid droplet due to altered regulation of ATGL; specifically, changes in the ratio of ATGL and PLIN5. We observed a reduced PLIN5/ATGL ratio in LD-enriched samples from HFD-fed mice compared to both Chow and HStD groups (Figure 5F), suggesting that ATGL has increased access to its lipid droplet- contained substrates [44]. From these observations we propose that insulin resistance is associated with dysregulated lipolysis in the liver, likely leading to poorly controlled fatty acid metabolic fluxes that spill over to bioactive lipid synthesis that influence hepatic insulin action [45, 46], which is not observed in insulin sensitive mice with fatty liver (Figure 5G).

## DISCUSSION

The liver plays a vital role in nutrient homeostasis, and lipid accumulation in the liver is linked to the development of many pathologies, including NAFLD/NASH, hepatocellular carcinoma (HCC), insulin resistance, and type 2 diabetes. Studies exploring the mechanisms linking fatty liver to metabolic dysregulation, including insulin resistance, have predominantly used HFDs compared to Chow or defined controlled diets. Using a novel dietary approach that induced three metabolic phenotypes – 1) lean, glucose tolerant, 2) obese, steatotic liver and glucose intolerant, and 3) obese, steatotic liver and glucose tolerant - we provide insights into the molecular events in lipid droplets of mice livers associated with lipid accumulation and those that associate with glucose intolerance and insulin resistance. From these observations, we propose that glucose intolerance and insulin resistance are associated with an imbalance in the lipolysome, consisting of lipolytic machinery [47], specifically the ratio of ATGL and PLIN5, which is sustained in insulin-sensitive mice with fatty liver. Overall, we provide evidence for the mechanisms that occur at the lipid droplets associated with the “safe” and “unsafe” storage of excessive lipids in the livers of mice.

The relationships between fatty liver disease, perturbed metabolic physiology and the development of pathologies such as type 2 diabetes, liver fibrosis and HCC are complicated. Like so many aspects of biology, these relationships are not binary, in that, not all patients with fatty liver progress to NASH, type 2 diabetes or HCC despite fatty liver being reported as a requisite for these conditions [5, 48, 49]. Epidemiological data clearly shows strong relationships between obesity and NAFLD - and the progression to NASH - regardless of metabolic health status [16, 50], while others have reported that metabolically healthy obese patients have less liver fibrosis compared to metabolically unhealthy obese patients [51]. Likewise, a recent meta-analysis reported that patients with obesity who are metabolically healthy without fatty liver had an increased risk of developing type 2 diabetes (pooled relative risk 1.42 (95%CI 1.11-1.77)), but that this was significantly less than that of patients with obesity who are metabolically healthy with fatty liver (pooled relative risk 3.28 (95%CI 2.30-4.67)), compared to metabolically healthy non-overweight subjects [52]. Studies have also reported that patients with obesity who are insulin sensitive have a low degree of liver steatosis compared to those who are obese and insulin resistant [27, 53], which is a pattern that persists in follow-up assessment [54]. The mechanisms that lead to fatty liver are many and include increased circulating fatty acid levels due to increased adiposity, increased *de novo* fatty acid synthesis and insufficient increase in mitochondrial fatty acid oxidation, as well as gene variants (see review [55]). Likewise, many mechanisms have been proposed to link fatty liver and metabolic dysfunction, including lipotoxic accumulation (including diacylglycerols and ceramides), oxidative stress, endoplasmic reticulum stress, impaired insulin signalling, and extrahepatic factors (see reviews [55–57]). In general, these proposed mechanisms arise from studies involving humans with or without fatty liver, or from rodent studies using modified diets such as HFD and high sucrose/fructose.

Our HStD provides a novel model to explore the relationships between obesity, adiposity, fatty liver and insulin resistance. Our detailed characterisation of this model, compared to Chow and HFD-fed mice, included assessment of adiposity, glucose tolerance, insulin sensitivity by hyperinsulinaemic-euglycaemic clamp, and tissue lipids by mass spectrometry [28]. In this study, we show that there was no relationship between lipid droplet morphology in the livers of HFD and HStD-fed mice and glucose intolerance. As such, mice fed HStD can safely store excess lipids in a manner that does not lead to metabolic dysregulation. This is analogous to the ‘Athlete’s Paradox’, where highly insulin-sensitive, endurance-trained athletes have skeletal muscle lipid levels similar to that observed in insulin-resistant obese and type 2 diabetes subjects [29]. Therefore, this approach provides a unique and powerful model to explore the relationships between obesity, fatty liver, and insulin action.

The accumulation of lipid droplets in hepatocytes is a hallmark feature of fatty liver. Lipid droplets serve as temporary storage sites for excess lipids, including fatty acids (stored as TGs), ceramides (stored as acyl-ceramides), and retinols and sterols (as esters) [58]. As such, the levels of lipids stored in lipid droplets, and thereby the size and number of lipid droplets, is the net effect of lipid synthesis and breakdown mechanisms that occur at the lipid droplet and the ER. Many studies have identified the lipid droplet proteome in rodent liver and in models of obesity and NAFLD [21–26]. However, these studies predominantly compare two groups, for example, between Chow (control, low-fat diet) and HFD-fed animals or fasting and fasting-re-fed states, and so identify changes in the lipid droplet proteomes that are associated with multiple differences between two groups, such as adiposity, tissue and circulating lipid levels, circulating hormone levels, and immune status. Including the HStD group that exhibits liver steatosis and glucose tolerance in our study design allowed us to identify changes in lipid droplet protein levels associated with fatty liver and those associated with glucose intolerance and insulin resistance. We identified 568 proteins (283 increased, 285 decreased) that were altered in fatty liver, but only 80 proteins (61 increased, 19 decreased) were associated with impaired glucose tolerance. It was unsurprising that fatty liver was associated with greater change in the lipid droplet proteome compared to glucose intolerance and insulin resistance. Many of the proteins that were enriched in the fatty liver proteome were involved in fatty acid metabolism and lipid droplet biology, including PLIN4 and MGLL, whereas FABP1 and ACBP_Dbi were less abundant. Consistent with Krahmer and colleagues [24], we observed changes in the levels of proteins involved in oxidative phosphorylation with HFD feeding, HStD feeding, and our fatty liver cluster. This suggests that increased lipid droplet number and size are associated with changes in inter-organelle contacts with mitochondria [35].

Loss and gain of function studies of key LD homeostasis regulators, such as DGATs [59, 60], ATGL [61, 62], ABHD5 [63], HILPDA/HIG2 [64], and PLIN5 [65], have provided significant insights into links between LD biology and liver and whole-body insulin action. However, these studies collectively demonstrate that there are complex relationships at play. In fact, both overexpression and knockdown of ATGL in mouse liver improved insulin action in a setting of HFD-induced insulin resistance [61, 62]. Combined with the multidimensional changes that occur with HFD feeding and associating changes in the LD proteome with insulin resistance, it has been challenging to uncouple steatosis-dependant changes in protein levels at liver lipid droplets from those associated with insulin resistance. In this study, the novel inclusion of the HStD group enabled the identification of 80 proteins whose abundance was associated with insulin resistance and glucose intolerance. Notable proteins identified included many involved in fatty acid metabolism and lipid droplet homeostasis, such as ABHD5, FABP5, ASCL4 and PLIN5. In the context of unchanged ATLG levels, the altered abundance of PLIN5 in LD is fascinating, as PLIN5 blocks ATGL-mediated lipolysis by competitively binding to ABHD5 and disrupting the interaction between ABHD5 and ATGL [44]. We observed a reduction in the ATGL to PLIN5 ratio in the insulin-resistant steatotic HFD group but not in the insulin-sensitive steatotic HStD and insulin- sensitive Chow groups, indicative of less PLIN5 to prevent ATGL TG hydrolase activity. As such, we predict that in the insulin-resistant setting, dysregulated lipolysis likely leads to increased availability of intracellular fatty acids to support bioactive lipid synthesis and impaired insulin action. This predicted increase in lipolytic activity may lead to reduced TG levels, and it is conceivable that the rates of synthesis and breakdown are matched to sustain TG levels. However, the flux of TG synthesis and breakdown differs between the HFD and HStD groups. PLIN5 plays roles in other aspects of fatty acid metabolism, which are regulated by the phosphorylation of Ser155 as well as being tissue/cell-specific [66]. *In vivo* quantification of hepatic lipolytic activity and PLIN5 function to validate this hypothesis is required but remains technically challenging to perform. Nonetheless, we provide novel insights into the proteomic changes at the LDs of livers of mice that associate with insulin resistance that are separate from changes due to increased TG levels.

Our findings demonstrate distinct changes in the liver LD proteome associated with the development of fatty liver that differ from those associated with insulin resistance. Furthermore, these changes occurred in settings with no LD number or size changes. Combined with our comprehensive metabolic characterisation of the HStD and HFD models [28], these data provide new insights into the complex relationships between the relationships between ectopic lipid accumulation, LD biology and insulin resistance.

## METHODS

### Animals

All surgical and experimental procedures performed were approved by the Animal Ethics Committee (University of Sydney) and were in accordance with the National Health and Medical Research Council of Australia’s guidelines on animal experimentation.

Eight-week-old male C57BL/6Arc mice were obtained from the Australian Animal Resource Centre (Perth, Australia). Mice were communally housed and maintained at 22±1 °C on a 12:12 hour light-dark cycle with ad libitum access to food and water with corn cob bedding.

Mice were assigned to one of three diets as previously described [28]. Briefly, the three diets were a standard Chow diet (11% fat, 23% protein and 66% carbohydrate, by calories; Specialty Feeds, Perth Australia), a high-fat diet (HFD; 60% fat, 20% carbohydrate (predominantly corn starch), 20% protein, by calories; based on Research Diets formula #D12492) or a high-starch diet (HStD; 20% protein, 20% fat, 60% carbohydrate (predominantly corn starch)) that were prepared in house, where the macronutrients of the diets contained identical amounts (in total grams) of commercially available vitamins (AIN vitamins) to that of the Chow diet. The energy density of the standard chow diet, the HFD and the HStD were 13 KJ/g, 13.4 KJ/g and 9.42 KJ/g, respectively. All cages were maintained on their assigned diets for 12 weeks. Food intake was performed at week 10 on mice by the daily weighing of food hoppers and food spillage in communally housed cages and was averaged to account for multiple mice per cage. Energy intake was calculated by multiplying the grams of food consumed with the energy density of the diet.

### Measurement of physiological parameters

Body composition was determined using the EchoMRI-500 (EchoMRI LLC, Houston, USA) according to the manufacturers’ instructions, excluding body water.

Glucose tolerance was determined in mice that were fasted for 6 hours (food removed at 8 am). After a basal sample, an oral bolus of 50 mg of glucose (200 µl of 25% glucose solution in water) was administered, and glucose levels were measured at 15, 30, 45, 60 and 90 mins post glucose load from tail vein blood using a glucose monitor (Accu-check Performa II, Roche Diagnostics, Australia). Insulin levels during the oral glucose tolerance test (Basal, 15 and 30 mins post load) were measured in samples of whole blood collected from the tail using a mouse ultra-sensitive ELISA kit (Crystal Chem, Elk Grove Village, USA).

### Analytical Methods

Plasma insulin was measured using the Ultra-Sensitive Mouse Insulin ELISA Kit from Crystal Chem, (USA). Liver triacylglycerols (TGs) were extracted using the method of Folch [67] and quantified using an enzymatic colourimetric method (GPO-PAP reagent; Roche Diagnostics).

### Lipid Droplet Isolation

Lipid droplet-enriched fractions were generated as previously published [68]. Briefly, fresh livers were diced and then homogenised in ice-cold hypotonic lysis medium (HLM; 20 mM Tris-Cl pH 7.4, 1 mM EDTA, 10 mM NaF supplemented with protease and phosphatase inhibitors (Astral Scientific)) at a ratio of 4 ml medium/ gram of tissue. Homogenates were then transferred to a 50 ml tube and centrifuged for 10 mins at 1 000 × *g* at 4°C (Beckman Coulter). The resulting supernatant was then transferred to microfuge tubes for storage at -80°C as whole liver lysates or to fresh ultracentrifuge tubes for further processing.

A sucrose gradient was prepared by mixing 3 ml of liver homogenate with 1 ml of HLM containing 60% sucrose in a 13.2 ml ultracentrifuge tube (Beckman Coulter), followed by 5 ml and then 4 ml of HLM buffer containing 5% and 0% sucrose, respectively. Samples were centrifuged for 30 mins at 28 000 × *g* at 4°C (P55ST2-636, Hitachi), with the rotor allowed to coast to a stop. The buoyant fraction (enriched with lipid droplets) was transferred to a 15 ml tube (Corning). Tubes containing the buoyant fraction were filled with 10 volumes of ice-cold acetone to de-lipidate the samples. Samples were incubated overnight at -20°C and centrifuged the next day for 1 hour at 4 300 × *g* at 4°C to pellet the proteins. A nitrogen sample concentrator was used to evaporate the residual acetone and dehydrate the pellet. Samples were then stored at -80°C until use.

### Immunoblotting

Whole liver lysates and lipid droplet-enriched fraction proteins were loaded on 10% SDS-PAGE gels and transferred onto polyvinylidene fluoride membranes (Merck). Membranes were incubated at room temperature for 1 h with blocking buffer (TBS, pH 4.5, with 0.1% Tween 20 (TBST) and 3% non-fat milk or BSA). Membranes were incubated overnight at 4°C with a specific primary antibody in TBST and 3% BSA. Antibodies used were as follows: lipid droplet - ATGL (1:1000, 2138S, CST) and PLIN2 (1:1000, ab108323, Abcam); mitochondria - Mitomix (1:500, ab110413, Abcam - Mitosciences) and cytochrome c (1:1000, 11940S, CST); cytosol - GAPDH (1:1000, 2118S/5174S, CST); golgi – GM130 (1:1000, 12480S, CST). Subsequently, membranes were incubated with secondary HRP-coupled antibodies (rabbit: 7074P2, mouse: 7076S. CST), washed again then incubated with enhanced chemiluminescence reagent (Merck) prior to visualisation using the ChemiDoc System (Bio-Rad Laboratories, Hercules, USA). Data were analysed using the ImageLab 5.2 version software (Bio-Rad Laboratories, Hercules, USA).

### Proteomic Analysis of Lipid Droplet Proteins

Lipid droplet proteins were resuspended in 4% sodium deoxycholate and 100 mM Tris-HCl (pH 7.5), and protein concentrations were quantified using a CBQCA Quantification kit (C-6667, Invitrogen) according to the manufacturer’s instructions. Samples were reduced using 10 mM TCEP and alkylated with 40 mM chloroacetamide at 95°C for 10 mins. Following this, samples were diluted to 1% sodium deoxycholate using Tris-HCl (pH 8) and digested overnight with MS-grade trypsin (in 50 mM acetic acid) at 37°C whilst constantly shaking. Peptides were submitted to sample clean-up as described previously [69], with the only change being that only the aqueous phase was put through the tips. Samples were dried for 1 h at 45°C in a centrifugal evaporator and stored in 5% (v/v) formic acid at 4°C before LC/MS-MS analysis.

These peptides were analysed using a Thermo Fisher Dionex RSLCnano UHPLC and directly added onto a 45 cm x 75 µm C-18 (Dr Maisch, Ammerbuch, Germany, 1.9 µm) fused silica analytical column with a 10 µm pulled tip, coupled to an online nano-spray ESI source. Peptides were resolved over a gradient from 5% ACN to 40% ACN running for 60 mins with a flow rate of 300 nL/min. Peptides were ionised by electrospray ionisation at 2.3 kV. Tandem mass spectrometry (MS/MS) analysis was performed using a Q-Exactive Plus mass spectrometer (Thermo Fisher) with higher-energy collisional dissociation fragmentation.

Data-dependent acquisition was used with the acquisition of MS/MS spectra for the top 10 most abundant ions at any one point during the gradient. Raw data were analysed using the quantitative proteomics software MaxQuant [70] (http://www.maxquant.org, version 1.5.7.0). Peptide and protein level identifications were both set to a false discovery rate of 1% using a target-decoy based strategy and proteins were filtered such that they had to have more than two razor and unique peptides. The database supplied to the search engine for peptide identifications contained the human UniProt database, downloaded on 30 September 2018, containing 42170 protein sequence entries and the MaxQuant contaminants database. Mass tolerance was set to 4.5 ppm for precursor ions and MS/MS mass tolerance was 20 ppm. Enzyme specificity was set to trypsin (cleavage C-terminal to Lys and Arg) with a maximum of 2 missed cleavages permitted. Deamidation of Asn and Gln, oxidation of Met, pyro-Glu (with peptide N-term Gln) and protein N-terminal acetylation were set as variable modifications. N-ethylmaleimide on Cys was searched as a fixed modification. The Max label-free quantification (LFQ) algorithm was used for LFQ, integrated into the MaxQuant environment [70, 71].

### Bioinformatic Analysis

Processing and statistical analysis of MaxQuant LFQ output was performed using the R software environment (version 3.4.3) as previously described [72]. For quantification, we applied a threshold that required an identified protein to be detected in at least seven of the ten mice in any of the three diets. Statistical outputs were corrected for multiple comparisons using the Benjamini-Hochberg method. Analysis of protein-protein interaction networks and functional enrichment was performed using the STRING database [73] and Kyoto Encyclopedia of Genes and Genomes (KEGG) pathways.

Significantly different lipid droplet-associated proteins were filtered to identify proteins of interest by creating biologically-relevant scenarios. Specifically, we identified proteins whose abundance correlated with liver TG levels using the following criteria: HFD vs Chow p ≤ 0.05, HStD vs Chow p ≤ 0.05, HStD vs HFD p ≥ 0.05. Next, we looked for proteins whose abundance correlated with glucose tolerance using the following criteria: HFD vs Chow p ≤ 0.05, Hi-ST vs Chow p ≥ 0.05, HStD vs HFD p ≤ 0.05.

### Liver Lipid Droplet Morphology

A portion of the liver was embedded in OCT and then cut into 10 µm sections using a cryotome FSE cryostat. A minimum of two cross-sections from different depths of the tissue was mounted on glass slide. Sections were covered with Oil Red O for 5 mins and thereafter carefully rinsed under running tap water for 30 mins. Next, slides were cover slipped with glycerol: water solution (9:1) and allowed to dry for 30 mins before sealing the edges with nail polish.

Sections were visualised with the Zeiss Axio Vert.A1microscope, and images captured with a Zeiss Axiocam 105 camera using the Zeiss software. Quantification of liver lipid droplet number and size was performed using ImageJ software (NIH, Bethesda, MA). For lipid droplet number, liver sections were divided into defined squares (0.145 mm^2^) and all lipid droplets within this area were counted for each mouse (n = 3 per group). For lipid droplet size, the same criteria were applied, and the average size for the total number of lipid droplets counted was calculated [74].

### Statistical Analysis

Statistical analyses were performed with GraphPad Prism 9.2.0 (GraphPad Software, San Diego, CA) or as described in Bioinformatic Analysis. Differences among groups were assessed with appropriate statistical tests noted in figure legends. *P* ≤ 0.05 was considered significant. Data are reported as mean ± SEM of at least 3 independent determinations.

Our interests were focused on identifying molecular links to phenotype, not the influence of diet per se. As such, we removed two HStD-fed mice from our analyses as they failed to respond to the diet, as determined by their body weight, adiposity and liver triacylglycerol (TG) levels (two SD away from the mean of the remaining mice, data not shown), and thereby failed to display the expected phenotype of the group. Additionally, one HFD-fed mouse was also removed from our analyses because of technical issues with the proteomics (Figure 1 - supplement S1A).

## AUTHOR CONTRIBUTIONS

*Andries Van Woerkom*: Investigation, Data curation, Formal analysis; *Dylan J Harney*: Investigation, Data curation, Formal analysis, Writing – review & editing; *Shilpa R. Nagarajan*: Investigation, Supervision; *Mariam F. Hakeem-Sanni*: Investigation; *Jinfeng Lin*: Investigation; *Matthew Hooke*: Investigation; *Tamara Pulpitel*: Investigation; *Gregory J Cooney*: Methodology, Writing – review & editing, Supervision; *Mark Larance*: Methodology, Resources, Software, Supervision; *Darren N. Saunders*: Data curation, Formal analysis, Software, Writing – review & editing, Visualization; *Amanda E Brandon*: Conceptualization, Formal analysis, Methodology, Writing – review & editing, Supervision, Project administration; *Andrew J. Hoy*: Conceptualization, Data curation, Formal analysis, Writing – original draft preparation, Writing – review & editing, Visualization, Supervision, Project administration, Funding acquisition.

## ACKNOWLEDGMENTS

AJH is supported by a Robinson Fellowship and funding from the University of Sydney. We thank Sydney Mass Spectrometry, a core research facility at the University of Sydney, for providing the Mass Spectrometry instruments used in this study.

**Supplementary Figure for Figure 3**: (A) Histograms showing peptide abundance distributions in individual replicates for each condition. (B) Venn diagram of identified proteins in each group. (C) Multiple regression analysis of peptide abundance in individual replicates. n=10 for Chow, n=9 for HFD, n=8 for HStD.

**Supplementary Figure for Figure 3. Immunoblots used as representative blots assembled in Figure 3**. All images were detected using ECL and Bio-Rad ChemiDoc System. Red boxes indicate the cropped portion of each immunoblot shown in the corresponding main figures.

**Supplementary Figure for Figure 5**: Abundance of known proteins that regulate lipid droplet biology in lipid droplet-enriched fractions of the liver of mice and the influence of modified diet feeding. Box-and-whisker plots for specific proteins of interest where each point represents the protein abundance in lipid droplet-enriched fractions for an individual mouse. n=10 for Chow, n=9 for HFD, n=8 for HStD. Data are presented as box and whisker plots: median, interquartile range and error bars representing Min to Max. * P≤0.05 vs. Chow; # *P*≤0.05 vs. HFD by t-test. LFQ, label-free quantification.

**Supplementary Table 1: LD LFQ Data Filtered**

**Supplementary Table 2: Chow LD proteome**

**Supplementary Table 3: HFD LD proteome**

**Supplementary Table 4: HStD LD proteome**

**Supplementary Table 5: Chow vs HFD proteins used for volcano plot**

**Supplementary Table 6: Chow vs HStD proteins used for volcano plot**

**Supplementary Table 7: HFD vs HStD proteins used for volcano plot**

**Supplementary Table 8: Differentially expressed proteome from ANOVA**

**Supplementary Table 9: Fatty liver scenario**

**Supplementary Table 10: Insulin sensitive scenario**

