## Supplementary figures and images for "Hepatic Lipid Droplet-Associated Proteome Changes Distinguish Dietary-Induced Fatty Liver from Insulin Resistance in Male Mice"

### Figure 3 - supplement S3

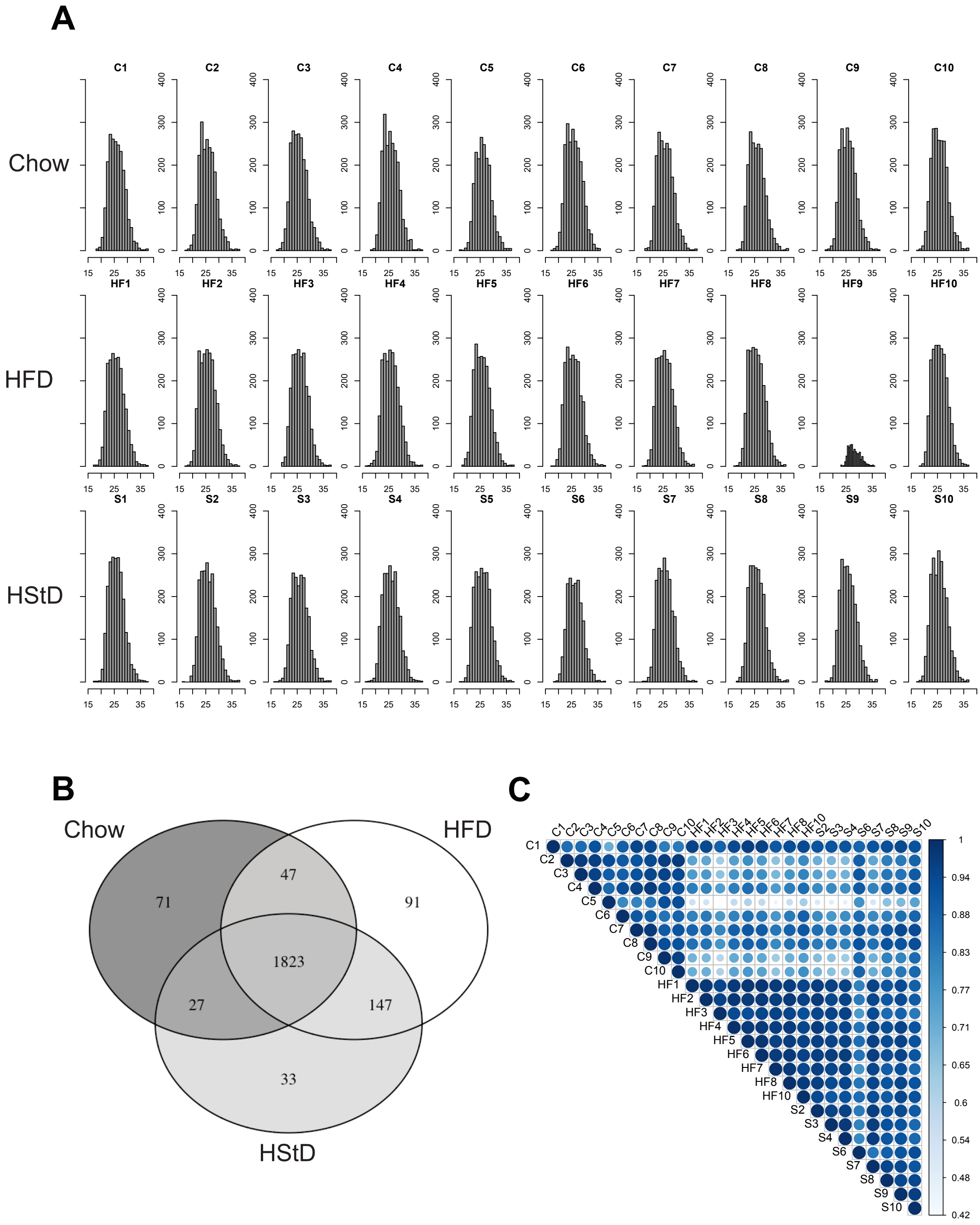

### Figure 5 - supplement S5

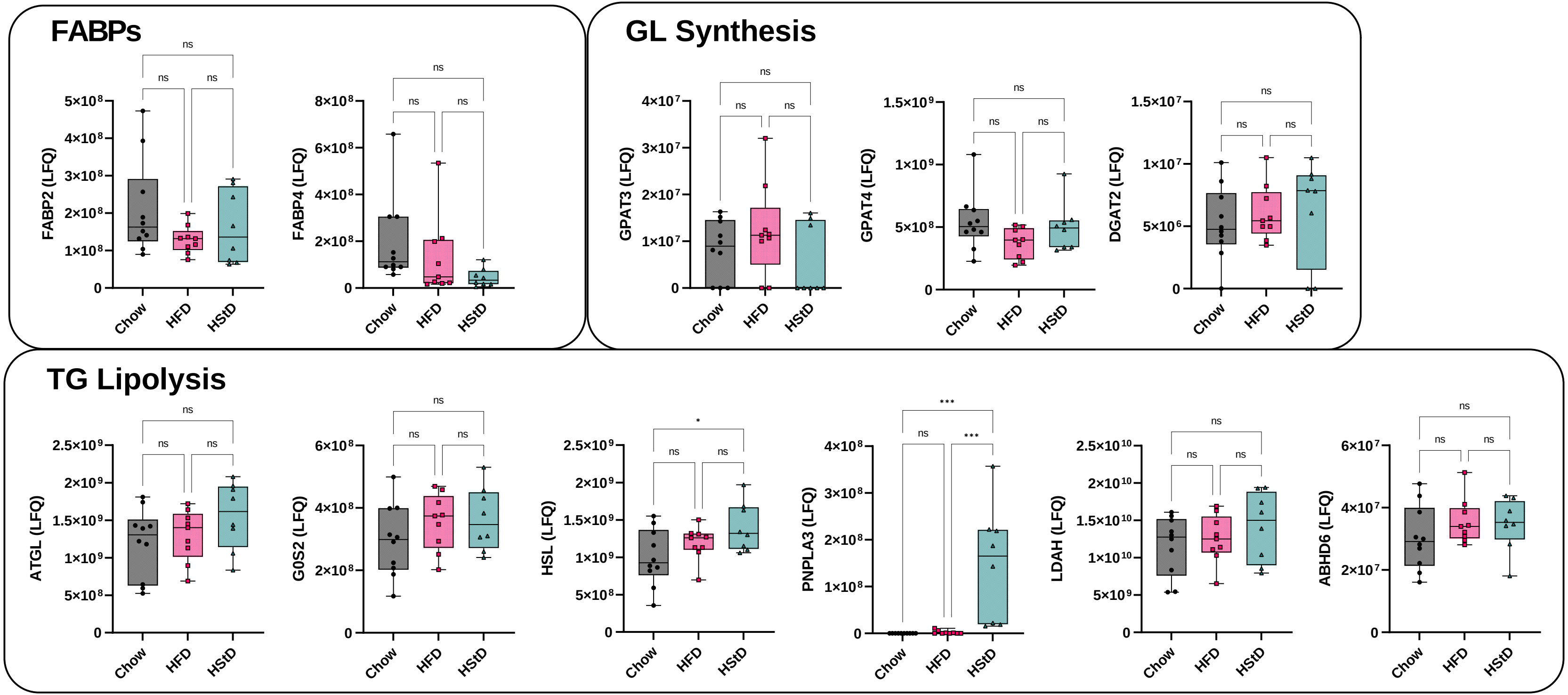
